# Synchronous nonmonotonic changes in functional connectivity and white matter integrity in a rat model of sporadic Alzheimer’s disease

**DOI:** 10.1101/2020.01.30.926444

**Authors:** Catarina Tristão Pereira, Yujian Diao, Ting Yin, Analina R da Silva, Bernard Lanz, Katarzyna Pierzchala, Carole Poitry-Yamate, Ileana O Jelescu

## Abstract

Brain glucose hypometabolism has been singled out as an important contributor and possibly main trigger to Alzheimer’s disease (AD). Intracerebroventricular injections of streptozotocin (icv-STZ) cause brain glucose hypometabolism without systemic diabetes. Here, a first-time longitudinal study of brain glucose metabolism, functional connectivity and white matter microstructure was performed in icv-STZ rats using PET and MRI. Histological markers of pathology were tested at an advanced stage of disease. STZ rats exhibited altered functional connectivity and intra-axonal damage and demyelination in brain regions typical of AD, in a temporal pattern of acute injury, transient recovery/compensation and chronic degeneration. In the context of sustained glucose hypometabolism, these nonmonotonic trends – also reported in behavioral studies of this animal model as well as in human AD – suggest a compensatory mechanism, possibly recruiting ketone bodies, that allows a partial and temporary repair of brain structure and function. The early acute phase could thus become a valuable therapeutic window to strengthen the recovery phase and prevent or delay chronic degeneration, to be considered both in preclinical and clinical studies of AD. In conclusion, this work reveals the consequences of brain insulin resistance on structure and function, highlights signature nonmonotonic trajectories in their evolution and proposes potent MRI-derived biomarkers translatable to human AD and diabetic populations.

## 1. Introduction

Alzheimer’s disease (AD) is the most common form of dementia and its incidence outburst already constitutes an economic and social burden. Clinically, AD is manifested by progressive memory loss and gradual decline in cognitive function, often culminating in premature death. However, pathological brain changes in AD begin to develop decades before the first symptoms (Sperling et al., 2011). The characterization of the temporal progression of these changes promises to provide an understanding of disease mechanisms, an effective disease staging and a window for therapeutic intervention.

In that regard, many studies have proposed the order in which relevant biomarkers occur in the progression of disease (Jack et al., 2013; Sperling et al., 2011). The pathological cascade of AD is characterized by initial deposition of amyloid-β (Aβ) plaques in the extracellular space and intracellular neurofibrillary tangles of hyperphosphorylated tau (NFTs) (Dickerson, 2013; Shaw et al., 2009; Wang et al., 2013). In addition, neurodegeneration manifests as reduced brain glucose metabolism (Edison et al., 2007), impaired synaptic activity (Shankar and Walsh, 2009) and neuronal loss (Attwell and Laughlin, 2001; Zarow et al., 2005). Gross cerebral atrophy (Gispert et al., 2015) and white matter degeneration (Chang et al., 2015; Choo et al., 2010; Dong et al., 2020; Jelescu et al., 2018) have been considered relevant biomarkers as well, along with reduced functional connectivity (Binnewijzend et al., 2012; Franzmeier et al., 2019). Nevertheless, the spatio-temporal relationship between AD hallmarks remains unclear.

Meanwhile, hypometabolism of glucose in the brain is taking the center stage as a key player in the onset of AD (Kuehn, 2020), while the amyloid cascade hypothesis is being questioned (Kametani and Hasegawa, 2018). Impaired brain insulin function has been reported to trigger AD pathology (Correia et al., 2011; Hölscher, 2019), while less robust glycolysis has been shown in animals at increased risk for developing dementia (Wu et al., 2018). Moreover, metabolic decline has been linked to impaired glucose transport and neuroinflammation (An et al., 2018; Balducci and Forloni, 2018). As it becomes well-established that diabetes makes the brain more susceptible to the aging process (Gudala et al., 2013), AD pathology has been suggested to be a brain-specific type-III diabetes (de la Monte and Wands, 2008).

Animal models allow studying each of the contributors to the cascade individually and obtaining comprehensive longitudinal data across their lifespan, which is particularly important for untangling direct effects of contributors and their interactions. Much effort has been put into the development and studies of transgenic mouse models of amyloidosis and tauopathy, which revealed for instance that – in mice – mutations causing amyloidosis alone do not produce NFTs, and tauopathy alone results in a fronto-temporal dementia pattern rather than AD (Buxbaum, 2009). The validity of transgenic animals is also limited in that they are not representative of the sporadic form of the disease, which is the predominant one in humans (King, 2018).

Numerous non-transgenic rat models have been proposed to replicate the AD phenotype, including models based on the ventricular infusion of the amyloid peptide (Sanganahalli et al., 2013); chemically-induced cholinergic-dysfunction models (Karthick et al., 2016); senescence-accelerated OXYS rats induced by a galactose-rich diet (Stefanova et al., 2015), and others (see (Benedikz et al., 2009) for a review). With the importance of brain glucose metabolism in AD being increasingly recognized, animal models of brain insulin resistance have been developed by an intracerebroventricular (icv) injection of streptozotocin (STZ). Intravenous injection of STZ is a very well-established model for type I diabetes (Ganda et al., 1976; Junod et al., 1969). Even though its mechanism is not fully elucidated, when delivered exclusively to the brain via the ventricles, this diabetogenic substance is thought to disrupt the brain insulin receptor system, thereby reducing glycolysis in the hippocampus and parieto-temporal cortex, but without causing systemic diabetes (Nitsch and Hoyer, 1991; Plaschke et al., 2010).

Accordingly, icv-STZ rats and monkeys have been shown to develop features typical of AD including memory impairment, NFT changes and extracellular accumulation of Aβ in the neocortex and hippocampus, thinning of the parietal cortex and corpus callosum (Knezovic et al., 2015), neuronal loss, astrocytosis and microgliosis in the corpus callosum and septum (Kraska et al., 2012), axonal damage and demyelination in the hippocampus and fimbria (Shoham et al., 2003) and decreased glucose uptake (Heo et al., 2011). Along with disruption of brain glucose metabolism, STZ induces oxidative stress by impairment of insulin signaling (Lester-Coll et al., 2006; Shoham et al., 2003).

In terms of STZ kinetics, previous studies have highlighted changes as early as one week and up to 39 weeks post-injection. Neurodegenerative lesions associated with inflammation have been reported in STZ rats at 1 and 13 weeks post-injection (Kraska et al., 2012). Tau and amyloid-β accumulation as well as cognitive deficits have been reported in STZ rats at 4, 13, 26 and 39 weeks post-injection (Knezovic et al., 2015). Finally, a FDG-PET study has shown glucose metabolism alterations in STZ monkeys at 6 and 12 weeks post-injection (Heo et al., 2011). The icv-STZ model has clearly demonstrated its potential to reproduce sporadic AD by the resemblance with the behavioral, neurochemical and structural changes found in the human AD brain. However, beyond anatomical data, this animal model has never been assessed with MRI, which could potentially offer significant insight into neurodegenerative processes.

We hypothesize that MRI-derived metrics of microstructure and functional connectivity in icv-STZ rats should show the same bi-phasic pattern as reported in human AD (Dickerson et al., 2005; Dong et al., 2020; Montal et al., 2018), with early inflammation followed by remission due to compensatory mechanisms and chronic neurodegeneration. Furthermore, in icv-STZ rats we could determine how these changes relate to glucose hypometabolism.

Here we therefore performed a longitudinal study in the icv-STZ rat model to quantitatively characterize alterations in functional connectivity and in white matter microstructure using resting-state functional (rs-fMRI) and diffusion MRI, respectively, as well as changes in brain glucose uptake captured by ^18^FDG (fluorodeoxyglucose)-PET. The diffusion MRI analysis was augmented with an extensively validated biophysical model of white matter which gives access to specific metrics of microstructure such as axonal density or intra-axonal diffusivity (Jelescu and Budde, 2017; Jespersen et al., 2018). Histological markers of pathology such as amyloid and NFTs were also tested. This study highlights the dynamics of how brain insulin resistance affects structure and function in a spatio-temporal pattern. It experimentally supports the caveat that metabolism underlies structure and function. This work further serves the dual purpose of outlining pathological trajectories relevant for clinical studies and validating potent MRI-derived biomarkers to track neurodegeneration in human AD and diabetes populations.

## 2. Methods

### 2.1. Animals

Animal experiments were approved by the Service for Veterinary Affairs of the canton of Vaud. Male Wistar rats (Charles River, l’Arbresle, France) weighing 218 - 259 g at baseline were assessed at three timepoints after the icv injection: PET and MRI data were collected at 2, 6 and13 weeks following surgery (**Figure 1**) and rats were sacrificed by perfusion fixation at 21 weeks for histological evaluation. Timepoints were chosen to be consistent with previous studies while also accommodation for constraints of repeated MRI and PET scanning related to anesthesia, cannulations, and scanner variability. Unfortunately, due to unforeseen technical issues with the PET scanner, the number of PET datasets was reduced at 2 weeks, and partially at 6 weeks. Later in the study, a major upgrade on the MRI system left the 13-week timepoint short of four MRI datasets.

**Figure 1.**
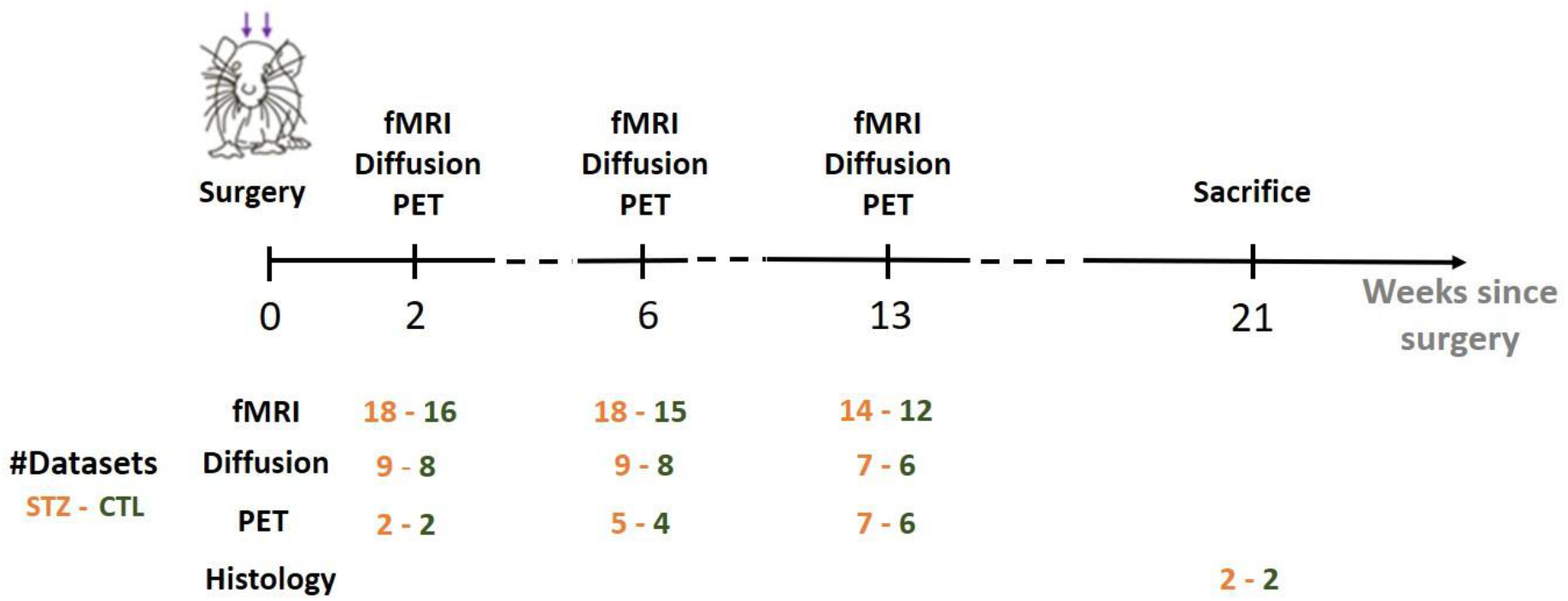
Experimental timeline. PET and MRI data were collected at 2, 6 and 13 weeks after icv-STZ injection. Two fMRI runs were acquired for each MRI session, which explains the larger number of datasets. Histology was performed after animal sacrifice at 21 weeks.

### 2.2. Intracerebroventricular injection

Rats were anesthetized using isoflurane (4% for induction and 2% for maintenance) and positioned in a stereotaxic frame (KopfModel900). The head was shaved and the skull was exposed with a midline incision. Two holes were burred into the skull, giving access to the two lateral ventricles at stereotaxic coordinates: ±1.4 mm lateral, 0.9 posterior and 3.5 mm deep relative to the bregma. A single bilateral icv injection (2 μL in each ventricle) of either 3 mg/kg STZ dissolved in citrate buffer (STZ group, n = 9), or citrate buffer solution (0.05 M, pH = 4.5) (control group, CTL, n = 8) was performed at a rate of 2 μL/min using Hamilton syringes, a double-syringe pump (TSE System) and catheters with 30G needles. The catheters were slowly removed 15 min after the end of the perfusion and the skin was sutured with silk sutures 3/0. 1500 mg paracetamol was administered in 800 mL drinking water starting 2 days before and up to 3 days after the surgery and an antiseptic cream was applied locally for 5 days after surgery. Rats were allowed to recover for two weeks before the first scanning session. They were kept in pairs in each cage for the entire study and checked daily by the veterinary team.

### 2.3. FDG-PET data acquisition

#### Animal preparation

Rats housed with free access to food and water were initially anesthetized using isoflurane (2% for induction) in an oxygen/air mixture of 30%/70% during tail vein cannulation for tracer delivery, and then isoflurane (1-2%) in 100% oxygen via a nose-cone once the animal was transferred and prone positioned on a temperature regulated PET scanner bed. Within the first minute of PET acquisition, a bolus of roughly 50 MBq ^18^F-FDG (Advanced Accelerator Applications, Geneva, Switzerland) in 50-300 μL was manually injected through the tail vein and followed by a saline chase. Throughout the experiment, breathing rate and body temperature were monitored using a respiration pillow and a rectal thermometer, respectively. Body temperature was maintained around (37±0.5) °C. Pre- and post-scan glycemia were determined from blood at the tail vein cannulation site using a Bayer glucose meter.

#### PET acquisition

All PET experiments were performed on an avalanche photodiode-based LabPET-4 small-animal scanner (Gamma Medica-Ideas Inc.) as described in (Lanz et al., 2014). Briefly, data were collected in list-mode and images between 30 and 50 minutes after injection were reconstructed using the maximum likelihood expectation maximization iterative reconstruction algorithm with a circular field of view (FOV) of 80 mm. The reconstructed voxel size was 0.25 x 0.25 x 1.18 mm. Steady-state radioactivity density images were then normalized for the effective injected FDG dose and the animal weight to generated Standardized Uptake Value (SUV) maps.

### 2.4. MRI data acquisition

#### Animal preparation

Animals were initially anesthetized using isoflurane (4% for induction and 1-2% for maintenance in an oxygen/air mixture of 30%/70%) and positioned in a homemade MRI cradle equipped with a fixation system (bite bar and ear bar). A catheter was inserted subcutaneously on the back of the animal for later medetomidine delivery. One hour before starting the rs-fMRI acquisition, anesthesia was switched from isoflurane to medetomidine (Dorbene, Graeub, Switzerland), which preserves neural activity and vascular response better than isoflurane (Pawela et al., 2009a; Weber et al., 2006), with an initial bolus of 0.1 mg/kg followed by a continuous perfusion of 0.1 mg/kg/h (Reynaud et al., 2019). The commercial solution at 1 mg/mL was diluted to 0.033 mg/mL. Throughout the experiment, the breathing rate and the body temperature were monitored similarly to the PET acquisition. The breathing rate under medetomidine was around 85 bpm. At the end of the scanning session, animals were woken up with an intramuscular injection of atipamezole (Alzane, Graeub, Switzerland) at 0.5 mg/kg.

#### MRI acquisition

All MRI experiments were performed on a 14 T Varian system (Abingdon, UK) equipped with 400 mT/m gradients. An in-house built quadrature surface coil was used for transmission and reception.

Structural T_2_-weighted images were collected using a fast spin-echo sequence with the following parameters: TE/TR = 10.17/3000 ms, echo train length: 4, matrix size = 128 x 128, FOV = 19.2 x 19.2 mm^2^, voxel size = 0.15 x 0.15 mm^2^, 30 coronal 0.5-mm slices, scan time = 10 minutes.

Diffusion-weighted data were acquired using a semi-adiabatic pulsed-gradient spin-echo (PGSE) segmented echo-planar-imaging (EPI) sequence (van de Looij et al., 2011), with the following protocol: 4 b = 0 images and 3 shells at b = 0.8 / 1.3 / 2.0 ms/μm^2^, with 12, 16 and 30 directions, respectively; δ/Δ = 4/27 ms; TE/TR = 48/2500 ms; 4 shots; matrix size = 128 x 64, FOV = 23 x 17 mm^2^, voxel size = 0.18 x 0.27 mm^2^, 9 coronal 1-mm slices, 4 repetitions, scan time = 1 hour.

Rs-fMRI data were acquired using a two-shot gradient-echo EPI sequence as follows: TE/TR = 10/800 ms, TRvol = 1.6 s, matrix size = 64 x 64, FOV = 23 x 23 mm^2^, voxel size = 0.36 x 0.36 mm^2^ and 8 coronal 1.12-mm slices, 370 repetitions, scan time = 10 minutes. Two fMRI runs were acquired in each MRI session.

### 2.5. Histology

At 21 weeks after the icv injection, rats were perfusion fixed using a 4% formaldehyde solution and the brains were promptly removed and immersed in 4% formaldehyde for two days, washed with PBS and stored at 4°C until they were embedded in paraffin and then cut on a sliding microtome into 8 μm thick slices. Mounted tissue sections were dried at room temperature overnight. Two brains per group were processed. Four main brain regions were investigated, centered around A-P atlas coordinates relative to bregma: +1mm and −0.4mm (covering the cingulate, motor and primary somatosensory (S1) cortices and the caudate putamen, −2.5mm (covering the retrosplenial (RSC) and S1 cortices, hippocampus, thalamus and hypothalamus) and −3.6 mm (covering the RSC, S1, auditory and parietal cortices, medial temporal lobe (MTL), hippocampus, thalamus and hypothalamus). Eight sections from each of these regions were stained with each of these contrasts: Congo red for amyloid-β plaques and Gallyas and Bielschowsky for NFTs.

#### Amyloid accumulation

Amyloid pathology was assessed with Congo Red staining (IHC WORLD, LLC). Slides were deparaffinized and rehydrated in distilled water, then incubated in Congo red working solution for 10 minutes and rinsed in distilled water. Finally, they were differentiated in alkaline alcohol solution, dehydrated through 95% and 100% alcohol then cleaned in xylene and mounted with resinous mounting medium.

#### Neurofibrillary tangles

Staging of tau protein pathology was first explored using the Gallyas Silver Stain method and the protocol described in https://www.protocolsonline.com/histology/dyes-and-stains/neurohistology/gallyas-silver-stain/.

#### Myelin staining

Myeloarchitecture was evaluated using LFB staining. Slides were deparaffinized and rehydrated, and then incubated in LFB solution at 56°C overnight in the dark. The excess stain was rinsed with 95% ethyl alcohol followed by distilled water. Then slides were differentiated in a lithium carbonate solution for 30 s, followed by 70% ethyl alcohol for 30 s, rinsed in distilled water, dehydrated in 100% alcohol, cleaned in xylene and finally mounted with resinous mounting medium.

#### Imaging

Stained slices were then imaged and digitized using a MEIJI TECHNO TC5600 Inverted Biological Microscope and were visually inspected for relevant staining features in STZ rats in comparison to the age-matched controls. Images were collected using the x5, x20 and x50 objectives at the same resolution.

### 2.6. Image processing

#### 2.6.1. FDG uptake

FDG-PET steady-state SUV maps were registered to their corresponding T2-weighted anatomical MR images with cross-correlation using ANTs (Avants et al., 2011, 2008), which was in turn registered to the Waxholm Space Atlas of the rat brain (https://www.nitrc.org/projects/whs-sd-atlas) using linear and non-linear registration (Avants et al., 2011) and 26 ROIs were automatically segmented. SUV images were normalized by the mean SUV over the brain to obtain SUVr maps corrected for inter-rat experimental variability (Heo et al., 2011).

#### 2.6.2. Resting-state functional connectivity

The rs-fMRI pre-processing pipeline is described in detail in (Diao et al., 2020). Briefly, images were skull-stripped, denoised (in the sense of thermal noise reduction) using a novel Marchenko Pastur Principle Component Analysis (MP-PCA) approach^1^ (Ades-Aron et al., 2018; Does et al., 2019; Veraart et al., 2016), corrected for EPI-related distortions using FSL’s topup (Smith et al., 2004), slice-timing corrected and spatially smoothed with a 0.36 x 0.36 x 1 mm^3^ full width half maximum (FWHM) kernel (Friston et al., 2007). Independent component analysis (ICA) was run on the fMRI timecourses using FSL’s MELODIC (Beckmann and Smith, 2004), with a high-pass filter (f > 0.01 Hz) and 40 components. All ICA decompositions were run through a dedicated rat FMRIB’s ICA-based Xnoiseifier (FIX) classifier (Diao et al., 2020), as previously described for humans and mice (Griffanti et al., 2017; Zerbi et al., 2015) and artefactual independent components attributable to physiological noise were automatically removed from the signal. Corrected fMRI images were registered to the anatomical image and then to the Waxholm Space Atlas using linear and non-linear registration in ANTs (Avants et al., 2008) and 28 atlas-defined ROIs (14 per hemisphere) were automatically segmented. Lastly, mean fMRI timecourses were extracted for each ROI based on “FIX-cleaned” data and individual ROI-to-ROI functional connectivity matrices were computed by calculating partial correlation between timecourses covarying for the global signal.

#### 2.6.3. Diffusion MRI and white matter microstructure

Pre-processing diffusion data included MP-PCA denoising (Veraart et al., 2016) followed by Gibbs-ringing correction^2^ (Kellner et al., 2016) and correction for EPI-related distortions, eddy currents and motion using FSL’s eddy (Andersson and Sotiropoulos, 2016). The diffusion and kurtosis tensors were estimated using a weighted linear least squares algorithm^3^ (Jensen et al., 2005; Veraart et al., 2013) and typical DTI and DKI-derived metrics were calculated: fractional anisotropy (FA), mean/axial/radial diffusivity (MD/AxD/RD) and mean/axial/radial kurtosis (MK/AK/RK). To overcome the lack of specificity inherent to empirical representations such as DTI and DKI, the biophysical WMTI model was computed in white matter voxels using a Watson distribution of axon orientations (Fieremans et al., 2011; Jespersen et al., 2018). The following microstructure parameters were extracted from the model: the intra-axonal water fraction *f,* the parallel intra-axonal diffusivity *D*_a_, the parallel *D*_e,║_ and perpendicular *D*_e,⊥_ extra-axonal diffusivities and the orientation distribution *C_2_* (**Figure 2**). This model has two mathematical solutions (Fieremans et al., 2011; Jelescu et al., 2016a; Novikov et al., 2018), of which the *D*_a_ > *D*_e,║_ solution was retained, according to recent evidence (Dhital et al., 2019; Jespersen et al., 2018; Kunz et al., 2018; Veraart et al., 2018). FA maps were registered to a FA template in the Waxholm Space using linear and non-linear registration in FSL (Jenkinson et al., 2002) and the corpus callosum, cingulum and fimbria of the hippocampus were automatically segmented. This approach enabled a more accurate segmentation of the white matter tracts, that otherwise tend to be misaligned due to enlargement of ventricles in STZ rats. Diffusion metrics were averaged over white matter ROIs.

**Figure 2:**
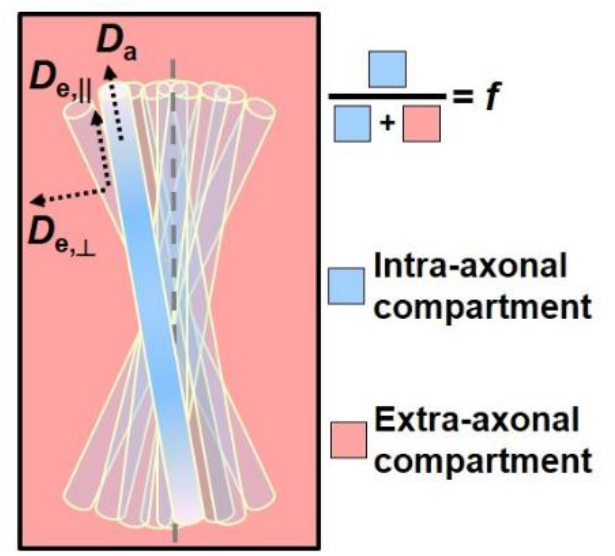
Schematics of the WMTI-Watson biophysical model (Jelescu and Budde, 2017; Jespersen et al., 2018). The diffusion signal is described in terms of two non-exchangeable compartments, the intra and extra-axonal spaces. Here, the axons are modelled as long narrow cylinders that reproduce a highly anisotropic medium. The intra-axonal space is described by a volume fraction of water molecules *f* and the parallel intra-axonal diffusivity *D*_a_. The perpendicular intra-axonal diffusivity is negligible at the relevant diffusion times, such that axons are considered to have radius equal to zero. The bundle of axons is embedded in the extra-axonal space that yields the parallel *D*_*e*,║_ and the perpendicular extra-axonal diffusivities *D*_*e*,⊥_. The axons’ orientations are modeled by a Watson distribution, which is characterized by 〈(*cosΨ*)^2^〉 ≡ *C*_2_.

### 2.7. Statistical analysis

The number of datasets available per group and timepoint in each type of analysis (PET, rs-fMRI and diffusion MRI) are reported in **Figure 1**.

#### FDG uptake

Regional differences in SUVr between STZ and CTL groups were evaluated at each timepoint (except at 2 weeks due to the small sample size) using one-tailed Mann-Whitney *U*-test (STZ < CTL), at significance level of α = 0.05. At 2 weeks, regional mean and standard deviation in each group were considered instead. The effect size of group differences was calculated using Hedge’s g corrected for small sample size at 6 and 13 weeks post-injection.

#### Resting-state functional connectivity

The significance of group differences was tested using non-parametric permutation tests (N=5000) with the Network-Based Statistics (NBS) Toolbox, with statistical significance retained at *p* = 0.05 after family-wise error rate (FWER) correction.

#### White matter microstructure

Average diffusion-derived metrics in each white matter ROI were compared between STZ and CTL groups at each timepoint using two-tailed Mann-Whitney *U*-test, at a α = 0.05 significance. Longitudinal changes within each group were assessed by one-way ANOVA and Tukey-Cramer correction, at a α = 0.05 significance. The effect size of group differences was calculated using corrected Hedge’s g.

#### Weight

Rats’ weight was compared between STZ and CTL groups at each timepoint using two-tailed Mann-Whitney *U*-test, at significance level of *α* = 0.05. In this analysis, we included 9/8 STZ/CTL rats at baseline, at 2 and 6 weeks after injection and 8/8 STZ/CTL rats at 13 weeks.

## 3. Results

Nine STZ rats and eight controls (CTL) were included in the longitudinal study. The datasets available for each modality (diffusion MRI, resting-state fMRI, FDG-PET and histology) at each of the four timepoints – 2, 6 and 13 weeks – are provided in **Figure 1**.

### 3.1. Weight

The weight of all animals was measured before the icv-STZ injection (baseline) and at all timepoints before each MRI or PET scanning session. There was no difference between groups at baseline, but STZ rats had a significantly lower weight than CTL rats at 2, 6 and 13 weeks after the injection (**Figure 3**).

**Figure 3:**
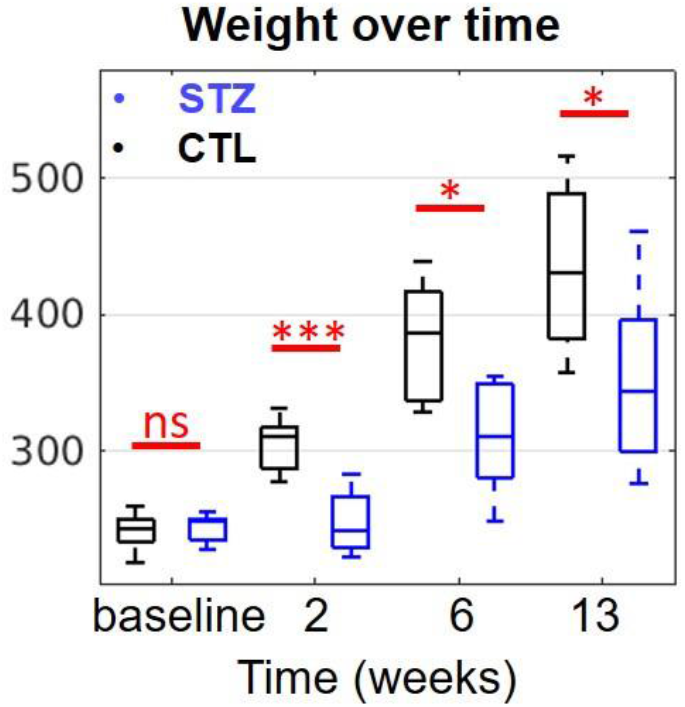
Group differences in animal weight measured before icv-STZ injection (baseline) and at all timepoints. Boxes represent median, 25^th^ and 75^th^ percentiles. ns: non-significant, *: *p* < 0.05, ***: *p* < 0.001 using the two-tailed Mann-Whitney *U*-test. 2 weeks: *p* = 0.0002, 6 weeks: *p* = 0.0152, 13 weeks: *p* = 0.0499.

### 3.2. FDG uptake

Representative SUVr maps are provided as **Supplementary Figure S1**. Relative Standardized Uptake Value (SUVr) was reduced in STZ rats compared to CTL rats, with different spatial patterns over time (**Figure 4**). At 2 weeks after injection, there were no differences between the STZ and the CTL groups in any of the regions of interest (ROIs) but the sample size was also very limited at that timepoint (2 animals per group). The most pronounced differences were found at 6 weeks after injection, involving several regions typically affected in AD. Lower SUVr was found in the anterior cingulate cortex (ACC) more significantly, but also in the retrosplenial cortex (RSC), the posterior parietal cortex (PPC), the motor, somatosensory, auditory and visual cortices. The medial temporal lobe (MTL) and the hippocampus (including subiculum) displayed a trend for lower SUVr with *p* < 0.1. Later, STZ rats showed less significant group differences throughout the brain, with the MTL being affected at 13 weeks. The effect sizes were large, varying between 1.16 and 2.06 (Hedge’s g values corrected for small sample size). **Supplementary Table S1** collects all *p*-values and corrected Hedge’s g values for group differences at each timepoint.

**Figure 4:**
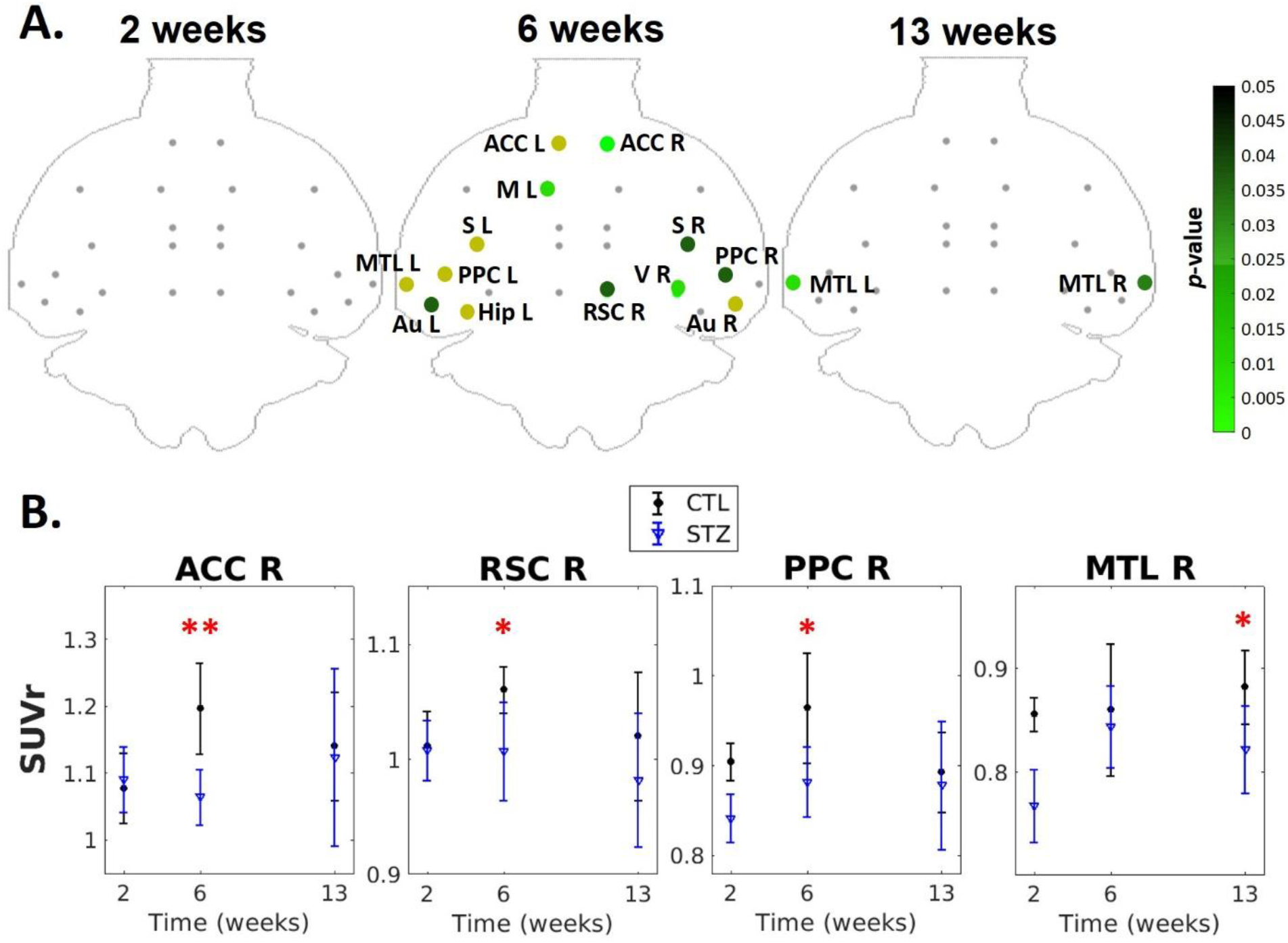
**A)** Group differences in SUVr at each timepoint. Green: ROIs with significantly lower SUV (p < 0.05 using one-tailed Mann-Whitney *U*-test, STZ < CTL). Dark yellow: Trend of lower SUV (p < 0.1). Correction for multiple comparisons was not applied given the small number of animals per group. *ACC: anterior cingulate cortex; RSC: retrosplenial cortex; PPC: posterior parietal cortex; MTL: medial temporal lobe; Hip: hippocampus; Au: auditory; V: visual; S: somatosensory and M: motor cortices; L/R: left/right.* **B)** Evolution of SUVr over time in four chosen ROIs of the right hemisphere. *\*: p < 0.05, **: p < 0.01.* The statistical analysis was not performed on the 2-week timepoint due to the small sample size. All p-values and corrected Hedge’s g values can be found in **Supplementary Table S1**.

### 3.3. Resting-state functional connectivity

Significant group differences in functional connectivity were found at 2, 6 and 13 weeks after injection (**Figure 5A**). In the **Figure 5B** graph networks highlight the group differences in nodal connections. At 2 weeks after injection, STZ rats displayed increased positive connectivity between the ACC and the RSC than CTL rats, and decreased anti-correlation of the default mode network (DMN) (including the ACC, RSC, hippocampus and subiculum) to regions of the lateral cortical network (LCN) including S1 and the motor cortex. At 6 weeks, the main changes in STZ rats remained reduced anti-correlations between the DMN (RSC, PPC and hippocampus) and LCN (S1/S2) as well as striatum. STZ rats also displayed increased connectivity between the hippocampus and the thalamus and between the MTL and the visual cortex compared to CTL rats. At 13 weeks, there was a dramatic shift towards overall reduced functional connectivity of the DMN in STZ rats, with weaker positive correlations between ACC, PPC, RSC, MTL, hippocampus, subiculum and visual cortex. STZ rats also presented a mild anti-correlation between the hypothalamus and the ACC, whereas these regions were uncorrelated in CTL rats. Remarkably, STZ rats exhibited a nonmonotonic trend in functional connectivity alterations, moving from hyperconnectivity within the DMN and less efficient dissociation between the DMN and LCN to hypoconnectivity within the DMN. This switch occurred between the 6 and 13-week timepoints.

**Figure 5:**
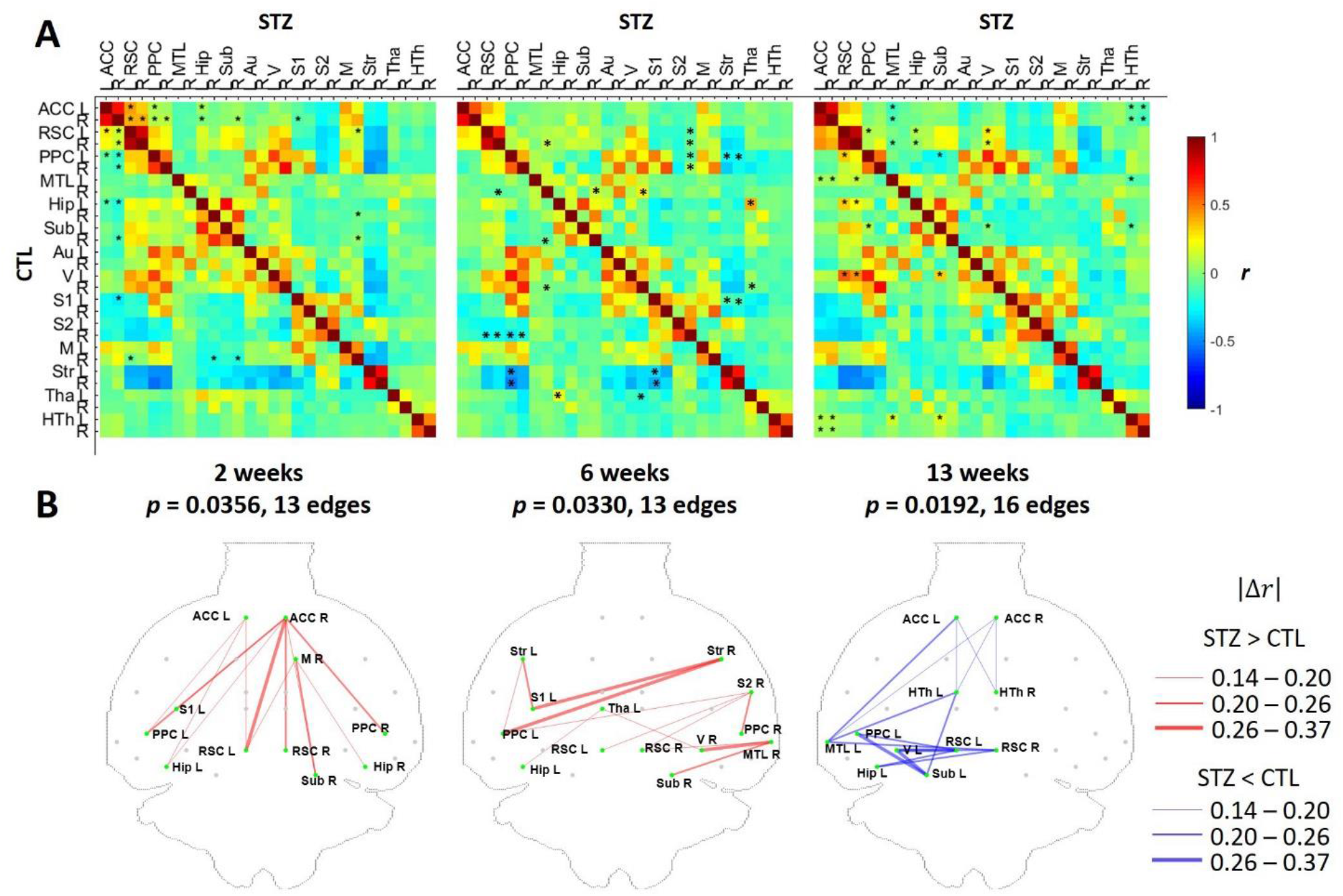
**A)** Hybrid average functional connectivity matrices at each timepoint (top-right half: STZ, bottom-left half: CTL). *: p < 0.05 (FWER corrected) at a threshold of 2.2. **B)** Graph networks at each timepoint. Red (STZ>CTL) or blue (STZ<CTL) edges and green nodes indicate connections with significant difference. Width of each connection represents the absolute difference in mean correlation coefficients |Δr|. The p-value for each network was given after FWER correction. *ACC: anterior cingulate cortex; RSC: retrosplenial cortex; PPC: posterior parietal cortex; MTL: medial temporal lobe; Hip: hippocampus; Sub: subiculum; Au: auditory; V: visual; S1/S2: primary/secondary somatosensory and M: motor cortices; Str: striatum; Tha: thalamus; HTh: hypothalamus; L/R: left/right.*

### 3.4. White matter microstructure

Representative diffusion images (*b* = 0 and FA maps) are provided in **Supplementary Figure S2**. Metrics derived from the classical Diffusion Tensor Imaging (DTI) technique provided differences between groups, which were very marked in the corpus callosum and fimbria of the hippocampus and milder in the cingulum (**Figure 6**). Compared to controls, STZ rats displayed lower fractional anisotropy (FA) in the corpus callosum at 2, 6 and 13 weeks after injection, in the fimbria at 2 and 13 weeks and in the cingulum at 13 weeks only. Metrics derived from Diffusion Kurtosis Imaging (DKI) – a clinically-feasible extension of DTI which also informs on tissue complexity – only captured group differences at 13 weeks after injection, when mean kurtosis (MK) was reduced in the STZ group in all ROIs. STZ rats also displayed lower axial (AK) and radial Kurtosis (RK) at 13 weeks in the corpus callosum and the fimbria and lower RK in the cingulum (not shown).. It is noteworthy that the fimbria of the hippocampus was the white matter tract with the largest differences between STZ and CTL rats.

**Figure 6:**
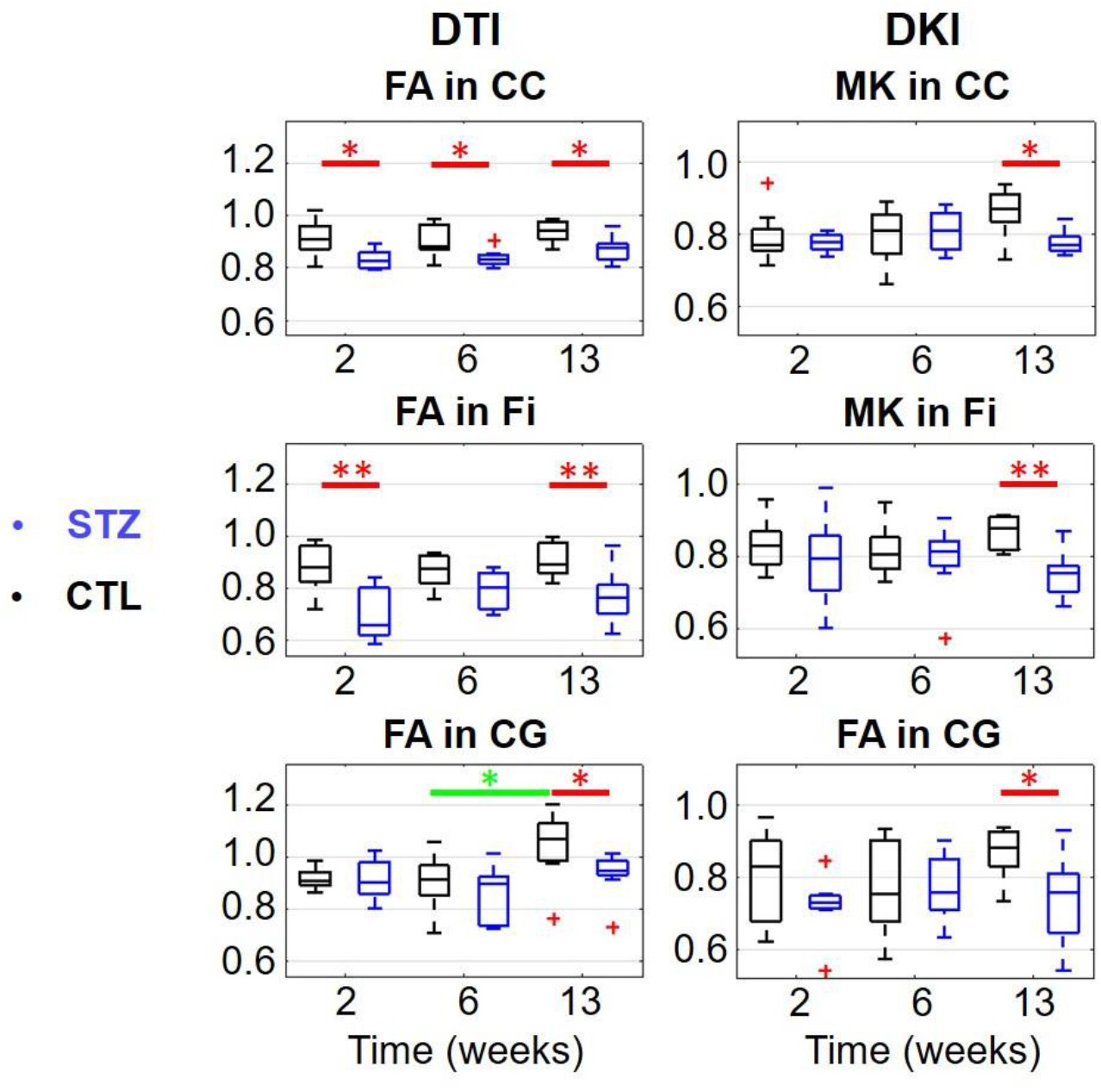
DTI and DKI estimates in 3 white matter ROIs (top row: corpus callosum (CC), middle row: fimbria of the hippocampus (Fi) and bottom row: cingulum (CG)). *FA: Fractional Anisotropy, MK: Mean Kurtosis.* Two-tailed Mann-Whitney *U*-test for inter-group comparison (red bars) and one-way ANOVA with Tukey-Cramer correction for within-group comparison across time (green bars). *: p < 0.05, **: p < 0.01. +: outlier values (but not excluded from the analysis). All p-values and corrected Hedge’s g values can be found in **Supplementary Table S2.**

The “White Matter Tract Integrity (WMTI)-Watson” biophysical model provided a more specific characterization of white matter degeneration than DTI and DKI. In the corpus callosum, both axonal fraction *f* and intra-axonal diffusivity *D_a_* were reduced in the STZ group at 2 weeks and 13 weeks (**Figure 7**). In the fimbria, *f* was lower in the STZ group at all timepoints, being more accentuated at 13 weeks, when *D_a_* was also lower. In the cingulum, reduced *f* was found at 13 weeks. Other model metrics did not show significant changes between groups or over time.

**Figure 7:**
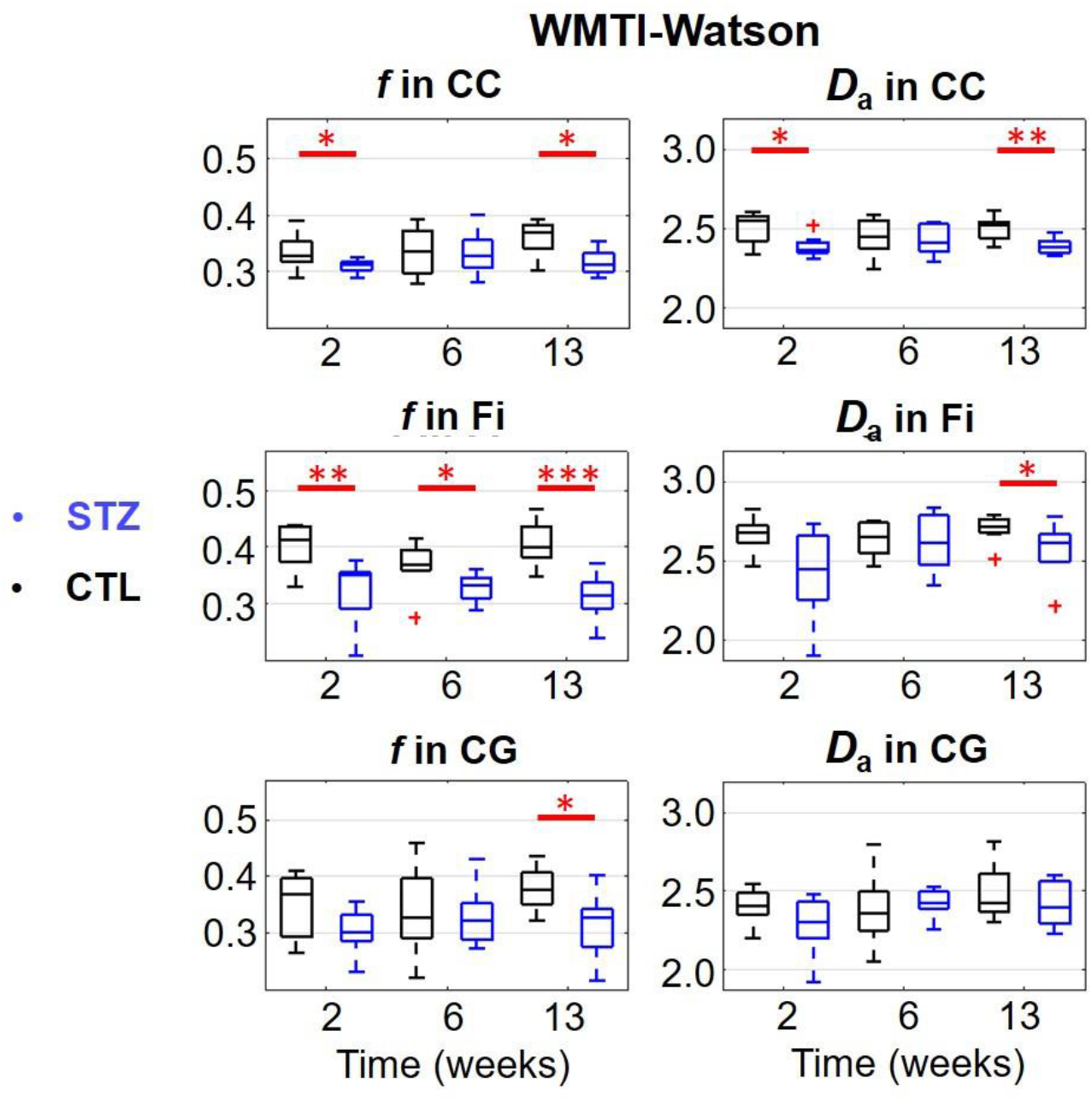
WMTI-Watson model estimates in 3 white matter ROIs (top row: corpus callosum (CC), middle row: fimbria of the hippocampus (Fi) and bottom row: cingulum *(CG)). f: axonal water fraction, D_a_: intra-axonal parallel diffusivity..* Two-tailed Mann-Whitney *U*-test for inter-group comparison (red bars).. *: p < 0.05, **: p < 0.01, ***: p < 0.001. +: outlier values (but not excluded from the analysis). All p-values and corrected Hedge’s g values can be found in **Supplementary Table S3.**

All group differences were supported by very large effect sizes, with absolute corrected Hedge’s g values varying between 0.84 and 1.83 in tensors derived metrics and between 0.87 and 2.12 in WMTI-derived metrics.

Longitudinal differences consisted in an increase in FA in the cingulum in the CTL between 6 and 13 weeks post-injection (*p* = 0.0348), while in the STZ group there was a significant decrease in AK in the corpus callosum between 6 and 13 weeks (*p* = 0.0357, not shown).

Group differences (*p*-values) and effect sizes (corrected Hedge’s g) at each timepoint for all tensor metrics and WMTI metrics are provided in Supplementary **Table S2** and **Table S3**, respectively.

Overall, white matter integrity displayed a nonmonotonic trend with partial recovery at 6 weeks, most markedly in the fimbria but also in the corpus callosum. The 13-week timepoint appeared as one of the most widespread damage, with involvement of the cingulum bundle as well.

### 3.5. Histology

At 21 weeks post-injection, corresponding to the study end-point, Congo red staining revealed amyloid accumulation, both intra and extra-cellularly, in the primary motor (M1), primary and secondary somatosensory (S1/S2, including the barrel field, trunk, dysgranular zone), agranular retrosplenial (RSA) and auditory (Au) cortices of STZ rats (**Figure 8**). In the motor and somatosensory cortex, an extracellular diffused red staining in the neighborhood of positively stained intracellular Aβ was observed (C, F), which can be related to diffused premature Aβ plaques. No amyloid deposits were found in other regions, and notably not in hippocampus.

**Figure 8.**
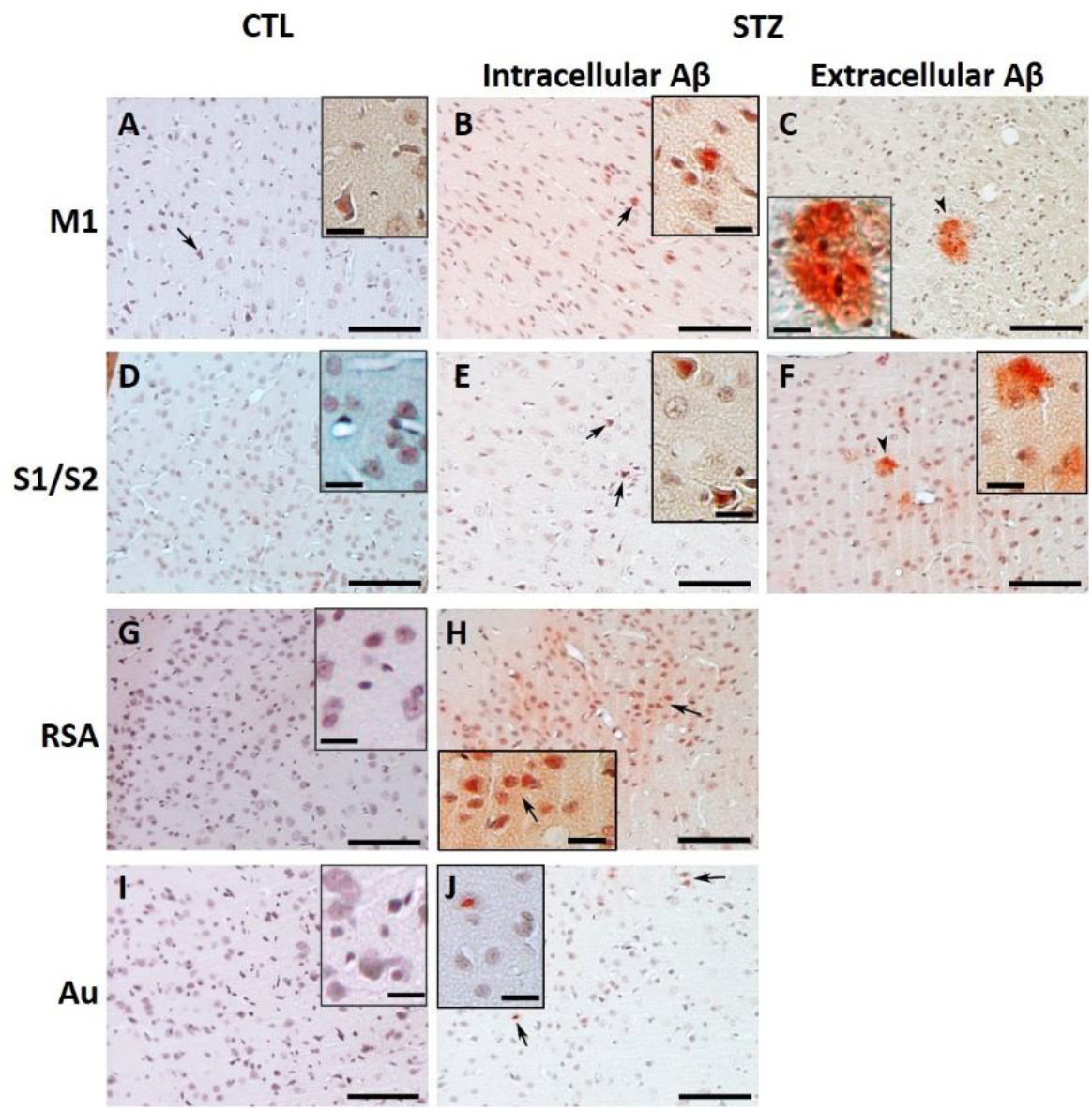
Aβ accumulation in the brain visualized with Congo red histochemical stain. Representative microphotographs of the brain of STZ rats (columns 2 and 3) and age-matched controls (column 1). The STZ treatment revealed positive signal of intracellular (column 2, arrows) and extracellular (column 3, arrowheads) Aβ accumulation in the primary motor (M1), primary and secondary somatosensory (S1/S2), agranular retrosplenial (RSA) and auditory (Au) cortices. Scale bar: 100 μm for main micrographs and 20 μm for insets.

Gallyas staining revealed abnormal tau-protein in the pre-tangle status in the cingulate (Cg1), primary motor (M1) and somatosensory cortices (S1) in both CTL and STZ rats (**Figure 9**). STZ animals also exhibited a more advanced form of NFTs in S1 (F). It should be noted that due to technical difficulties, Gallyas stain was compromised on the slices corresponding to more posterior brain regions (AP < 0 mm), precluding the assessment of NFTs in posterior cortical areas and hippocampus.

**Figure 9.**
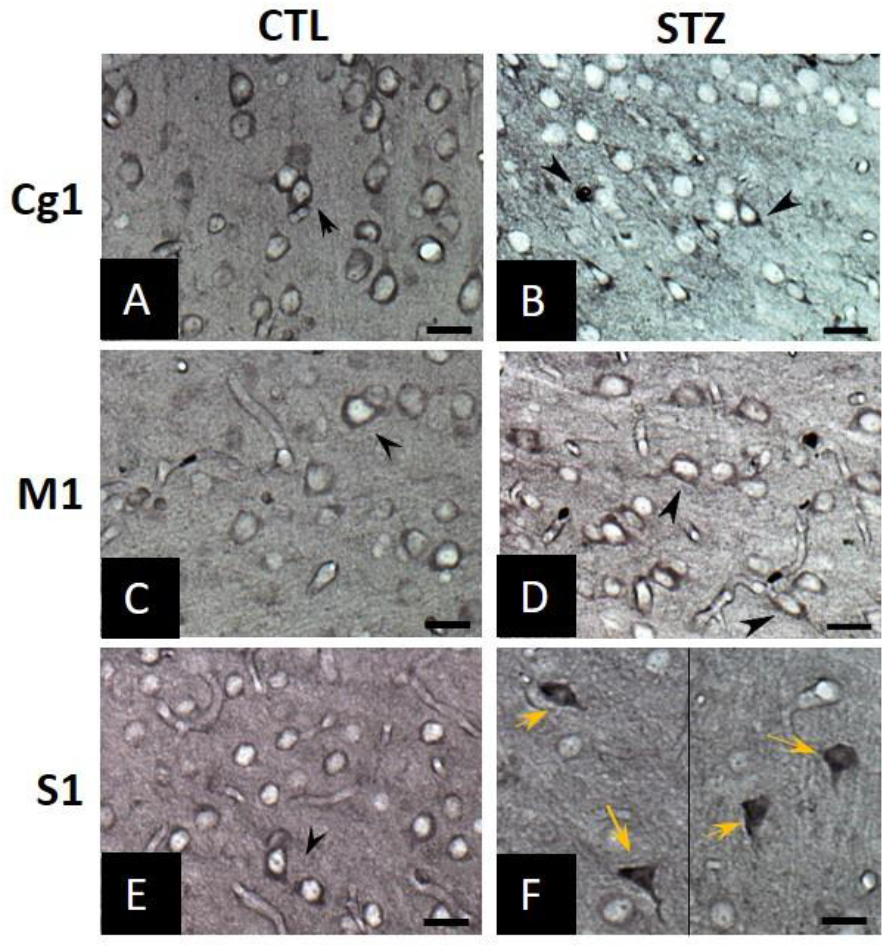
Representative micrographs of Gallyas silver stain of STZ rats (B, D, F) and age-matched controls (A, C, E) revealed accumulation of abnormal tau-protein in the pre-tangle status (black arrowheads) in the cingulate cortex (Cg1), primary motor (M1) and somatosensory cortex (S1) in both groups. The STZ animals exhibited however more advanced forms of NFTs in S1 (yellow arrows). Scale bar: 20 μm.

Luxol fast blue (LFB) revealed macroscopic atrophy of brain structures, particularly the fimbria, and enlargement of the lateral ventricles in STZ rats. This feature was also apparent on *in vivo* MRI anatomical images (**Figure 10**).

**Figure 10.**
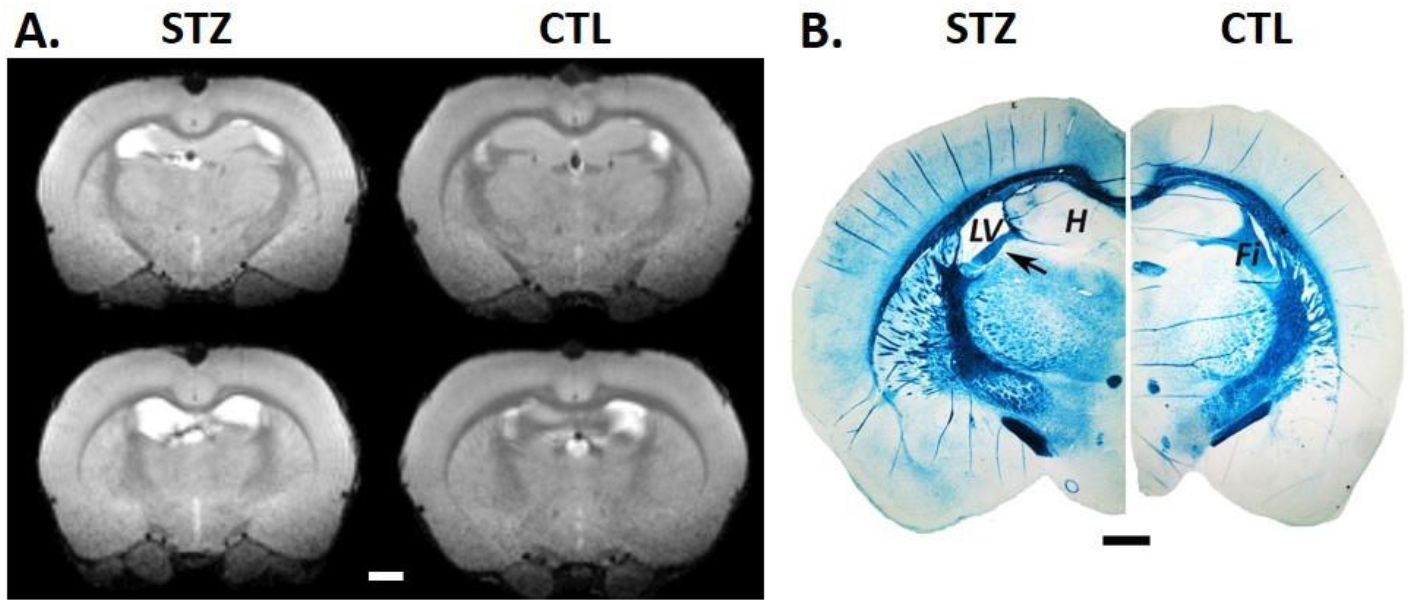
Atrophy of the fimbria (Fi) and enlargement of lateral ventricle (LV) in STZ rats vs CTL at 21 weeks after icv injection. **A)** *In vivo* MRI anatomical image in two axial planes (anterior (top row) to posterior (bottom row) position). Scale bar: 2 mm. **B)** LFB staining, confirming shrinkage of the fimbria (arrow) and significant enlargement of ventricular space (LV) in STZ rats. Scale bar 1 mm. *H: hippocampus.*

**Figure 11:**
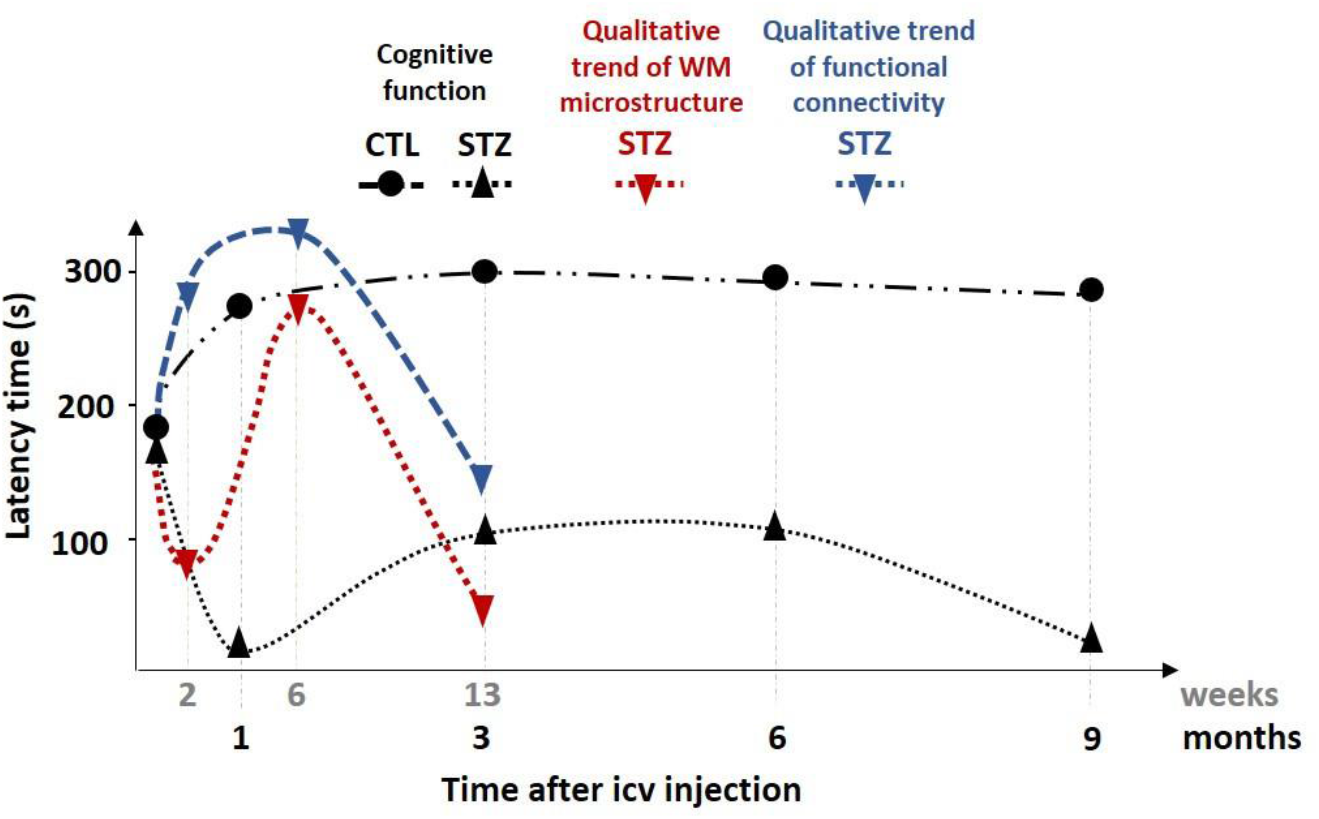
Progression of MRI biomarkers in comparison with memory performance in the icv-STZ rat model. Figure adapted from (Knezovic et al., 2015). Black: Latency time measured in the passive avoidance test reported in (Knezovic et al., 2015) as a function of time in months.Red/blue: Qualitative trend of the diffusion/functional MRI biomarkers – respectively - reported in the present study as function of time in weeks. The plots of cognitive performance in CTL and STZ rats correspond to the follow-up of male Wistar rats that received an initial injection of either 3 mg/kg of STZ or buffer only, same as in our experiment. Differences between the procedures in Knezovic et al. and our experiments lie in the weight of rats at baseline (300 g vs 250 g, respectively) and the fact that the STZ dose was split over 2 injections (day 1 and day 3) vs single injection, respectively. The timepoints of observation were also not identical but the dynamics are slower in the memory performance, with a chronic stage occurring around 26 weeks, while MRI-derived biomarkers point to an earlier switch towards pronounced neurodegeneration and reduced functional connectivity between 6 and 13 weeks.

## 4. Discussion and conclusions

The icv-STZ animal model has reportedly shown alterations typical of AD, reinforcing the idea that neurodegeneration might be primarily induced by an insulin resistance in the brain. Its effects on the pathological cascade are however not fully understood, and the present study attempts to characterize it by means of MRI-derived biomarkers for the first time. In this longitudinal assessment, FDG uptake, resting-state functional connectivity and white matter microstructure results agree in terms of spatio-temporal variations and are consistent with our histological findings and with previous studies of cognitive decline and immunochemistry.

Functional connectivity in the CTL group was consistent with previous reports that midline regions of the DMN are anti-correlated with the sensorimotor system (Gozzi and Schwarz, 2016). In STZ animals, these anti-correlations between DMN and LCN were initially reduced (2 and 6 weeks) suggesting less efficient network dissociation and brain processing (Fox et al., 2005). Concomitantly, hyperconnectivity within the DMN was found in STZ rats at these early timepoints, broadly consistent with initial hyperconnectivity of the hippocampus in human studies of mild cognitive impairment (Dickerson et al., 2005). By 13 weeks however, STZ rats exhibited on the contrary reduced connectivity (or hypoconnectivity) in regions typically involved in AD, also consistent with eventual memory impairment in this animal model and reduced functional connectivity in AD patients (Binnewijzend et al., 2012; Franzmeier et al., 2019). Changes in functional connectivity thus displayed a nonmonotonic pattern, with initial hyper-connectivity and impaired network dissociation, followed by later hypo-connectivity. The switch occurred between the 6 and 13-week timepoints.

White matter degeneration is best assessed when examining results from the DTI/DKI analysis and from the WMTI-Watson model jointly. In the corpus callosum, reduced anisotropy was detected throughout the 2-13 week span, but subtle specific differences, in the form of reduced axonal fraction *f* and intra-axonal diffusivity *D*_a_ for example, were only found at 2 and 13 weeks, not at 6 weeks. Similarly, reduced anisotropy in the fimbria was detected at 2 and 13 weeks, while the WMTI-Watson model revealed reduced axonal fraction *f* throughout the 2-13 week span, but with least pronounced differences at 6 weeks. Thus the 6-week timepoint appears as one of partial recovery, before a marked degeneration at 13 week, with reduced anisotropy, mean kurtosis, axonal fraction and intra-axonal diffusivity in all three bundles examined.

DTI and DKI metrics are indicative of a loss of tissue structure and complexity but do not provide specific information on the underlying changes. The characterization of white matter damage was further improved by biophysical modeling. Reduced intra-axonal diffusivity *D_a_* and axonal fraction *f* were indicative of acute intra-axonal injury and demyelination at 2 weeks, followed by partial remission at 6 weeks and finally chronic injury and demyelination, potentially leading up to axonal loss at 13 weeks. While demyelination can lead to increased extra-axonal diffusivity perpendicularly to axons (*D*_e,⊥_), our previous work using electron microscopy validation has shown that the demyelination needs to be widespread for this parameter to be altered, while “patchy” demyelination will immediately be manifest as decreased axonal fraction *f* (myelin is invisible in diffusion MRI and lost myelin is replaced by extra-axonal space) (Jelescu et al., 2016b). The absence of changes in the other model parameters (water diffusivity in the extra-axonal space, or axons orientation dispersion) contributes to a specific picture of white matter microstructure alterations, with most damage being attributed to the axonal space, which is consistent with intra-cellular toxic action of STZ (Knezovic et al., 2015; Lester-Coll et al., 2006). Histological evidence of neuronal loss leading to macroscopic thinning of corpus callosum (Knezovic et al., 2015) and of axonal damage and demyelination of the fimbria (Shoham et al., 2003) have been previously reported in STZ rats and attributed to oxidative stress (Shoham et al., 2003). In humans, partial loss of myelin, axons and oligodendroglial cells have been observed histologically in early AD (Brun and Englund, 1986; Englund et al., 1988; Kobayashi et al., 2002; Roher et al., 2002; Svennerholm and Gottfries, 1994). Recent non-invasive MRI methods including diffusion MRI and relaxometry also provide evidence for demyelination in mild cognitive impairment and early AD (Bouhrara et al., 2018; Dong et al., 2020). Current theories further suggest a two-step demyelination process, with initial oligodendrocyte degeneration releasing iron, which promotes amyloid plaques, which in turn further destroy myelin until the neurodegenerative stage (Bartzokis, 2011, 2004).

Overall, changes in functional connectivity agreed with those in white matter microstructure both spatially and temporally. Remarkably, hypo-connectivity of cingulate cortex and hippocampus at 13 weeks coincided with altered axonal integrity and myelination of the cingulum bundle appearing at that timepoint. Temporally, the initial acute damage and partial recovery in white matter microstructure at 2 and 6 weeks corresponds to an early phase of hyper-connectivity and impaired network dissociation, suggesting compensatory mechanisms, while the 13-week timepoint reveals widespread and pronounced white matter damage as well as loss of functional connectivity as markers of a chronic phase of disease.

Post-mortem histology revealed pathological accumulation of amyloid in STZ rats in motor, somatosensory, retrosplenial and auditory cortices. Abnormal tau was also found in anterior cingulate, motor and somatosensory areas, though NFTs specific to the STZ group were found in S1 only and information from posterior parts of the brain was unfortunately missing. Furthermore, our histology measurements are only available at a later timepoint (21 weeks after icv injection). Histology reports at multiple timepoints on this animal model are available in the literature (Knezovic et al., 2015; Kraska et al., 2012; Shoham et al., 2003). While enlarged ventricles are not specific to AD (Apostolova et al., 2012), here we observed more substantial brain atrophy and enlarged ventricles in STZ animals as compared to controls in both the in vivo MRI data and postmortem histology, which were fully consistent with previous reports (Kraska et al., 2012; Shoham et al., 2003).

Our MRI findings of axonal integrity and functional connectivity were consistent with reported neuronal loss in the septum and corpus callosum (Kraska et al., 2012), thinning of the parietal cortex and the corpus callosum (Knezovic et al., 2015) and demyelination associated with microglia activation in the hippocampus and fimbria (Shoham et al., 2003). However, our histology analysis at 21 weeks found no amyloid in the hippocampus and this region could unfortunately not be evaluated for NFTs, while early neurofibrillary changes in the hippocampus have been reported at 12 weeks followed by presence of diffuse Aβ plaques at 26 weeks (Knezovic et al., 2015).

PET data showed reduced FDG uptake in icv-STZ rats in brain regions typically involved in AD, and were consistent with reported changes in FDG uptake in the medial temporal lobe of icv-STZ monkeys as early as 2 weeks after injection (Heo et al., 2011). In our study, the most pronounced differences in uptake were found at 6 weeks, but the development of earlier glucose hypometabolism in STZ rats is highly likely. Indeed, the absence of SUVr group differences at the earliest timepoint was most probably due to a very limited PET cohort size (2 animals per group, **Figure 1**), with STZ rats nonetheless displaying lower SUVr in PPC and MTL, qualitatively. Furthermore, STZ rats were on average already 70 g lighter than CTL rats at that timepoint (**Figure 3**). The significant weight difference supports the assumption of impaired brain glucose metabolism as early as 2 weeks and the use of alternative energy sources such as stored body fat, which when burnt results in a buildup of ketones. Ketone bodies have been referred to as a “brain reserve” that can be mobilized to compensate for an inability to efficiently utilize glucose (Gano et al., 2014; Wu et al., 2018). It is remarkable that early weight loss has also been reported in AD (Sergi et al., 2013; Tamura et al., 2008), although various explanations for it have been proposed, including a low leptin state associated with hypothalamic dysfunction (Ishii et al., 2014). Interestingly, STZ rats showed increased anti-correlations between the ACC and the hypothalamus connectivity at 13 weeks after injection. Along these lines, we propose that weight loss and/or slower weight gain in our STZ rats reflect: (1) a change in energy homeostasis; and (2) the effect of ketones on the signalling in the hypothalamus of appetite regulating neurohormones, such as leptin, that respond to changes in carbohydrate (glucose/glycogen) to fat macronutrient availability (Sumithran et al., 2013; Thio, 2012). Remarkably, we report “partial recovery” in terms of axonal integrity and functional hyperconnectivity at six weeks post-injection, in a context of marked FDG uptake differences at that timepoint. This observation follows the caveat that metabolism underlies structure and function, and supports the assumption that past the acute phase, the brain switches to other forms of energy, such as ketone bodies, as a means to maintain energy homeostasis and restore normal brain function (Yang et al., 2019). Interestingly, the same regions with reduced FDG uptake at 6 weeks after STZ injection eventually showed functional hypoconnectivity at 13 weeks (**Figure 4** and **Figure 5**), supporting the manifestation of a chronic phase of disease. Here, our data suggests that neurodegeneration resumes as pathological mechanisms approach a critical stage and can no longer be compensated for.

The succession of acute change, compensation/partial remission and chronic degeneration phases, which exemplifies nonmonotonic changes, has also been reported in a nine-month follow-up of memory performance in icv-STZ rats, measured using a passive avoidance test (Knezovic et al., 2015). In this cognitive study however, the examination timepoints were different from ours and the acute phase was found at 1 month after injection, the recovery period between 3 and 6 months and chronic decrease of latency time from 6 months on. The curves of progression of the MRI-derived biomarkers of neurodegeneration used in this study were qualitatively overlaid with that of memory performance from (Knezovic et al., 2015) for comparison (Error! Reference source not found.). The qualitative trends in white matter microstructure and functional connectivity provide potentially earlier evidence of brain damage in the icv-STZ model than behavioral observations, as MRI-derived biomarkers are expected to be more sensitive to mechanisms occurring in the preclinical stage of the pathological cascade. Such neuroimaging trends could potentially predict the following symptomatic stage of the disease that includes cognitive deficits.

Nonmonotonic and biphasic trajectories of brain alterations have also been reported in several human studies of AD and mild cognitive impairment, and involve a variety of biomarkers from cortical volumes, cerebral perfusion, gray and white matter microstructure, and hippocampal activity (Dickerson et al., 2005; Dong et al., 2020; Fortea et al., 2014, 2011; Montal et al., 2018; Pegueroles et al., 2017; Sierra-Marcos, 2017), which reinforces the suitability of the icv-STZ animal model for sporadic AD. Furthermore, the non-monotonic progression of AD in humans was reported based on cross-sectional studies including subjects at different stages of disease. The current study in icv-STZ rats highlights this nonmonotonic pattern in a longitudinal framework also, which is easier to achieve in animals than in humans. The acute phase identified using MRI-derived biomarkers could represent a valuable therapeutic window to strengthen the recovery and prevent or delay chronic degeneration, to be considered both in preclinical and clinical studies of AD.

The quantification of functional connectivity and axonal degeneration has provided insightful information about spatial and temporal AD-like changes induced by brain glucose hypometabolism. The present study contributes to the growing hypothesis that brain insulin resistance is a key player in the onset of AD and the icv-STZ rat model offers great potential for the investigation of neurodegeneration in the context of AD and diabetes. For a complete picture of the pathological cascade in this animal model, a future longitudinal study should include MRI, PET, behavior and cross-sectional histology all at once.

### Study limitations

PET and fMRI experiments were conducted under different anaesthetic conditions, which may complicate the comparison of brain activity between the two modalities, but not the between-group comparisons for each modality. Isoflurane induces a very stable and controlled anaesthesia, which is well-suited for the safety requirements on PET experiments. The value of medetomidine vs isoflurane for rs-fMRI studies has however been largely shown in the literature (Kalthoff et al., 2013; Kint et al., 2020; Pawela et al., 2009b), which is why this sedative was chosen for fMRI. Most recent literature suggests that a combination of medetomidine and low-dose isoflurane may be ideal for rat fMRI (Kint et al., 2020; Paasonen et al., 2018) and this is something that will be considered in future studies.

As AD is more prevalent in women than men, studies in animal models should ideally characterize both sexes to assess the full representativeness of the model. However, as hormonal modulation seems to play an important role in the vulnerability or resistance to AD progression, female rats should ideally be included at an advanced aged, as an equivalent of women’s menopause. For example, a recent study has shown that hippocampal changes in icv-STZ rats were dependent on sex, remarkably with young female rats more resistant to STZ-induced alterations than males (Biasibetti et al., 2017). It is however not clear whether older females would not be more vulnerable to icv-STZ as a result of substantial hormonal changes. In our study, we used male rats only, which we acknowledge as a limitation. This choice was guided by practical reasons. However, we do plan to perform a similar study in aging female rats in the near future. More studies are indeed needed to better understand how gender differences and hormonal modulation can impact disease progression.

### Summary

We used the icv-STZ rat model to characterize longitudinal alterations in brain functional connectivity and white matter integrity resulting from impaired brain glucose metabolism. Icv-STZ animals displayed early hyper-connectivity within DMN and impaired network dissociation between DMN and LCN followed by hypo-connectivity within DMN.. Moreover, white matter degeneration was manifested as intra-axonal damage and demyelination, potentially leading to axonal loss and mainly affecting the corpus callosum and the fimbria of the hippocampus. It followed a sequence of early acute injury followed by transient partial recovery and late chronic degeneration. Structural and functional changes presented matching spatio-temporal patterns, with a turning point between 6 and 13 weeks, and were in agreement with existing literature. Histology at a later timepoint confirmed pathological accumulation of amyloid and NFTs in STZ rats, in gray matter regions also impacted by altered functional connectivity. In parallel, brain glucose hypometabolism was likely established from the start. The initial hyper-connectivity and transient repair in brain structure at six weeks post-STZ injection is interpreted as the recruitment of alternative forms of energy such as ketone bodies to make up for impaired glucose metabolism, until pathological mechanisms are no longer reversible. Our results support the hypothesis of impaired glucose metabolism as a key player in the onset of AD and the relevance of the icv-STZ animal models for pre- clinical AD research. This study (i) characterized the impact of brain glucose hypometabolism on brain microstructure and function, (ii), highlighted signature nonmonotonic trajectories in their evolution and (iii) proposed potent MRI-derived biomarkers translatable to human AD and diabetic populations to monitor disease progression and potential response to treatment.

## Acknowledgments

The authors thank Stefan Mitrea, Mario Lepore and Dario Sessa for technical assistance and the Histology Core Facility of the EPFL for staining procedures. This work was supported by the Centre d’Imagerie BioMédicale (CIBM) of the University of Lausanne (UNIL), the Swiss Federal Institute of Technology of Lausanne (EPFL), the University of Geneva (UniGe), the Centre Hospitalier Universitaire Vaudois (CHUV) and the Hôpitaux Universitaires de Genève (HUG). C.T.P. acknowledges the support of the Swiss-European Mobility Plan (Movetia).

## Supplementary Material

**Figure S1:**
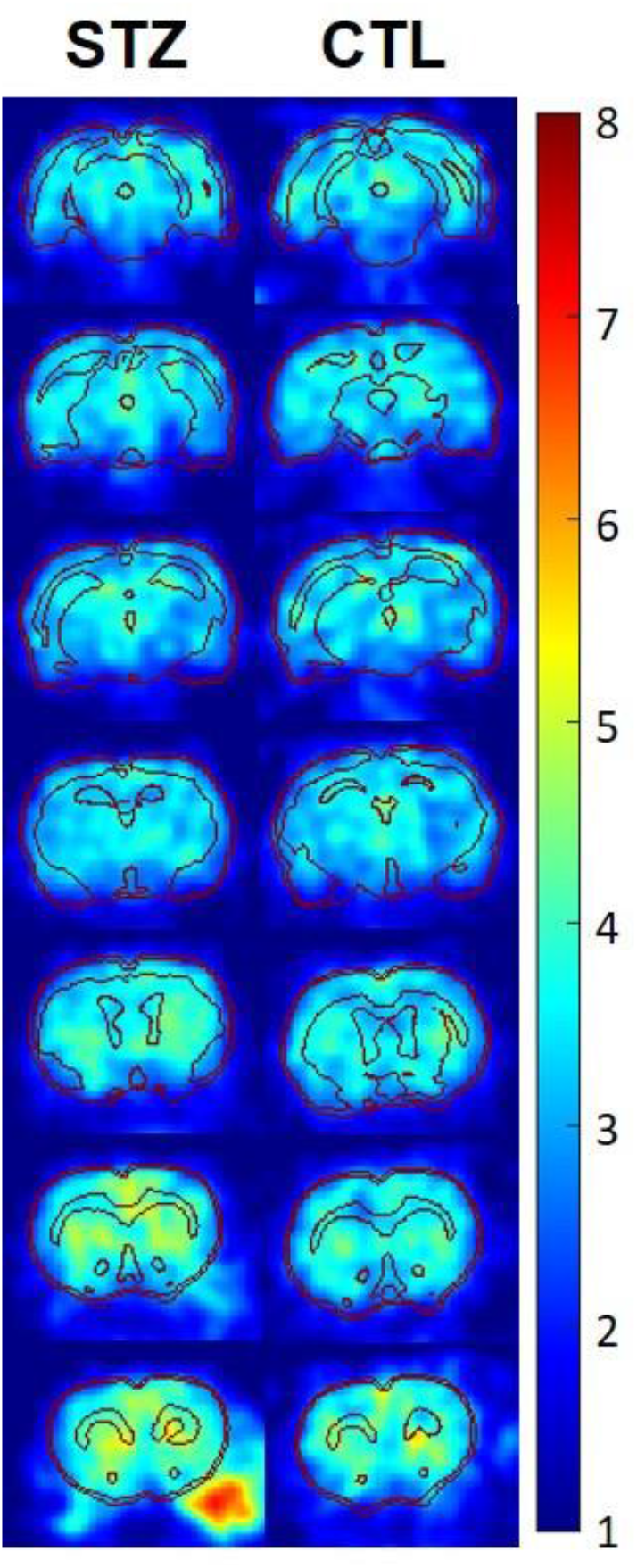
Multiplanar coronal SUV maps of one STZ and one CTL rat at 13 weeks after injection. MRI-based segmentation is overlaid onto the PET image for anatomical reference. Top to bottom: posterior to anterior.

**Figure S2:**
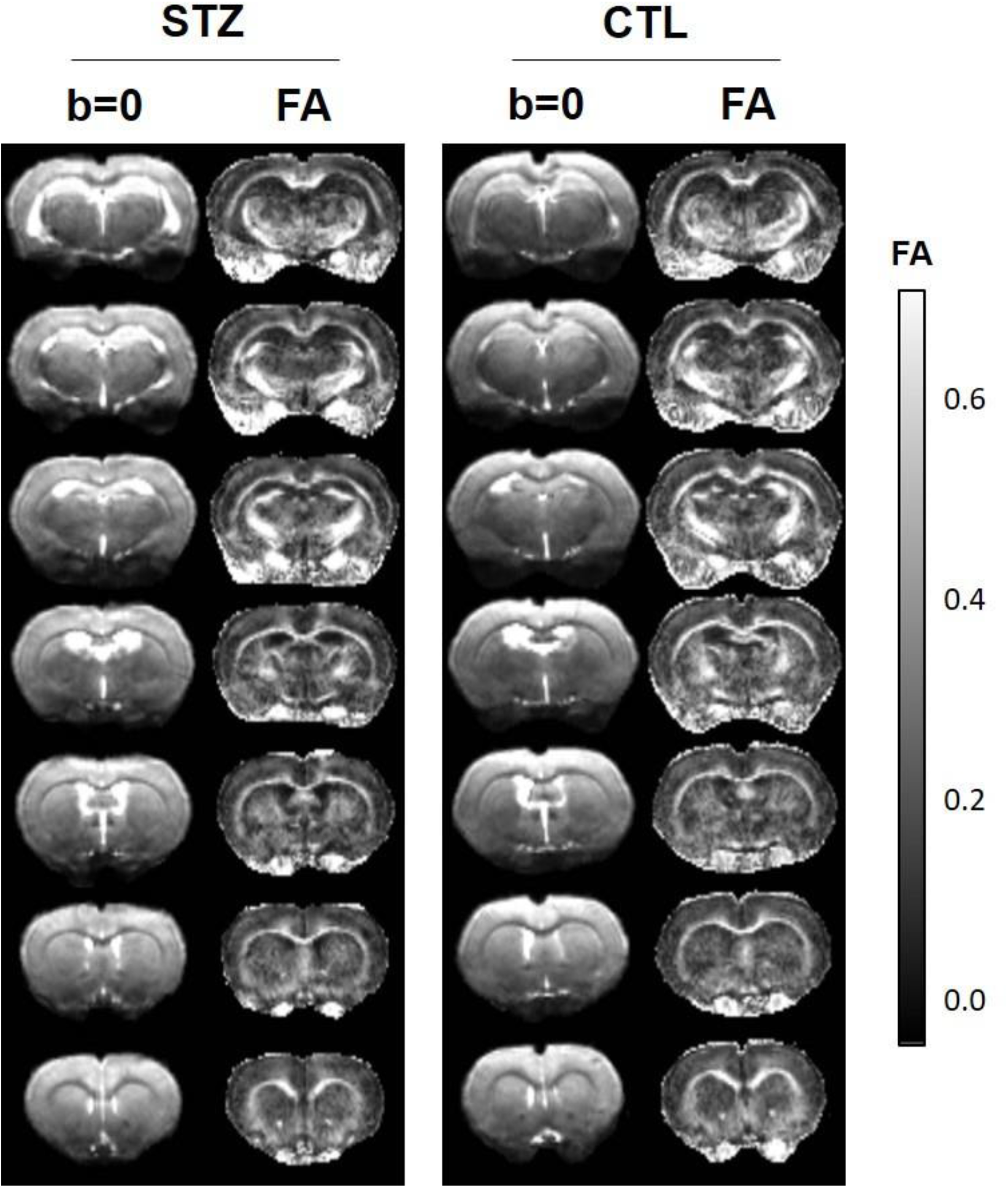
Multiplanar coronal b=0 images (T2 weighted, no diffusion weighting, arbitrary units) and fractional anisotropy (FA) maps of one STZ and one CTL rat at 13 weeks after injection. Top to bottom: posterior to anterior.

**Table S1:** List of p-values obtained from the one-tailed Mann-Whiteney U-test (α=0.05) and corrected Hedge’s g (g_corr_) values used to evaluate group differences in SUVr at each timepoint in each ROI. *Bold font: p < 0.05. Gray cells: ROIs shown in Figure 4.*

| Group differences in SUVr |  |  |  |  |  |  |
| --- | --- | --- | --- | --- | --- | --- |
| ROI | Time after injection |  |  |  |  |  |
|  | 2 weeks |  | 6 weeks |  | 13 weeks |  |
| | p-value | $g_{corr}$ | p-value | $g_{corr}$ | p-value | $g_{corr}$ |
| ACC L | - | - | 0.0556 | 1.2390 | 0.1830 | 0.3016 |
| ACC R | - | - | **0.0079** | **2.0641** | 0.2226 | 0.1305 |
| RSC L | - | - | 0.1429 | 0.7269 | 0.2669 | 0.2882 |
| RSC R | - | - | **0.0317** | **1.3380** | 0.1474 | 0.5778 |
| PPC L | - | - | 0.0556 | 0.9618 | 0.3654 | 0.0419 |
| PPC R | - | - | **0.0317** | **1.3933** | 0.3654 | 0.2104 |
| MTL L | - | - | 0.0556 | 1.1997 | **0.0111** | **1.2110** |
| MTL R | - | - | 0.3651 | 0.2713 | **0.0173** | **1.3112** |
| Hip L | - | - | 0.0556 | 1.1505 | 0.6346 | -0.1876 |
| Hip R | - | - | 0.2063 | 0.5389 | 0.7774 | -0.4618 |
| Au L | - | - | **0.0317** | **1.1603** | 0.2669 | 0.3879 |
| Au R | - | - | 0.0952 | 1.1500 | 0.1171 | 0.6268 |
| V L | - | - | 0.1429 | 0.7640 | 0.3141 | 0.1543 |
| V R | - | - | **0.0159** | **1.8793** | 0.1171 | 0.3583 |
| S1 L | - | - | 0.0952 | 1.0171 | 0.3141 | 0.2070 |
| S1 R | - | - | **0.0317** | **1.3407** | 0.1830 | 0.6583 |
| M L | - | - | **0.0079** | **1.6376** | 0.2226 | 0.4134 |
| M R | - | - | 0.2063 | 0.6322 | 0.1474 | 0.5432 |
| Str L | - | - | 0.3651 | 0.3583 | 0.4178 | -0.0562 |
| Str R | - | - | 0.3651 | 0.2498 | 0.3654 | -0.0170 |
| Tha L | - | - | 0.4524 | 0.1220 | 0.8829 | -0.4842 |
| Tha R | - | - | 0.7222 | -0.3358 | 0.8829 | -0.5244 |
| HTh L | - | - | 0.5476 | -0.1346 | 0.1830 | 0.4402 |
| HTh R | - | - | 0.9444 | -0.9164 | 0.6859 | -0.1436 |

**Table S2:** List of p-values obtained from the two-tailed Mann-Whitney U-test (α=0.05) and corrected Hedge’s g (g_corr_) used to evaluate differences in diffusion tensor (DTI and DKI)-derived metrics between STZ and CTL groups at each timepoint in each ROI. ROI: Region of interest; CC: corpus callosum; CG: cingulum; Fi: fimbria; FA: Fractional Anisotropy; MD: Mean Diffusivity; AD: Axial Diffusivity; RD: Radial Diffusivity; MK: Mean Kurtosis; AK: Axial Kurtosis; RK: Radial Kurtosis. *Bold font: p < 0.05. Gray cells: ROIs shown in Figure 6.*

| Group differences in tensors-derived metrics |  |  |  |  |  |  |  |
| --- | --- | --- | --- | --- | --- | --- | --- |
| DTI and DKI metrics | ROI | Time after injection (weeks) |  |  |  |  |  |
|  |  | 2 |  | 6 |  | 13 |  |
| | | p-value | $g_{corr}$ | p-value | $g_{corr}$ | p-value | $g_{corr}$ |
| FA | CC | **0.0104** | **1.3134** | **0.0281** | **1.1880** | **0.0111** | **1.3451** |
|  | CG | 0.8785 | 0.0230 | 0.5737 | 0.3946 | **0.0274** | **0.8274** |
|  | Fi | **0.0030** | **1.6863** | 0.0830 | 0.9270 | **0.0025** | **1.4660** |
| MD | CC | 0.5054 | -0.2442 | 0.2345 | -0.4247 | 0.9626 | 0.0713 |
|  | CG | 0.0650 | 1.0035 | 0.7209 | -0.0423 | 0.5414 | 0.2496 |
|  | Fi | 0.6454 | -0.0308 | 0.8785 | 0.0454 | 0.0927 | -0.8965 |
| AD | CC | 0.0830 | 0.8410 | 1.0000 | 0.1295 | 0.1672 | 0.6714 |
|  | CG | 0.0650 | 0.9984 | 0.8785 | 0.1219 | 0.0927 | 0.8123 |
|  | Fi | 0.7209 | 0.4950 | 0.7984 | 0.2983 | 1.0000 | -0.0812 |
| RD | CC | 0.1304 | -0.8765 | 0.1049 | -0.7536 | 0.1672 | -0.4644 |
|  | CG | 0.1304 | 0.8474 | 0.6454 | -0.1336 | 0.5414 | -0.3136 |
|  | Fi | 0.1949 | -0.5435 | 0.5054 | -0.1953 | **0.0206** | **-1.3531** |
| MK | CC | 0.8785 | 0.2632 | 0.9591 | -0.1768 | **0.0152** | **1.4166** |
|  | CG | 0.1949 | 0.6536 | 1.0000 | 0.0014 | **0.0206** | **1.1457** |
|  | Fi | 0.4418 | 0.3735 | 0.9591 | 0.2437 | **0.0016** | **1.8393** |
| AK | CC | 1.0000 | 0.1886 | 0.3823 | -0.4796 | **0.0079** | **1.5719** |
|  | CG | 0.6454 | 0.3835 | 0.7209 | -0.1894 | 0.2359 | 0.5698 |
|  | Fi | 0.7984 | -0.0856 | 0.5737 | -0.0365 | **0.0464** | **1.1686** |
| RK | CC | 0.8785 | 0.2859 | 0.9591 | 0.0782 | **0.0274** | **1.1341** |
|  | CG | 0.2345 | 0.8498 | 0.8785 | 0.1682 | **0.0152** | **1.2823** |
|  | Fi | 0.2345 | 0.7142 | 0.0830 | 1.0204 | **0.0079** | **1.3696** |

**Table S3:**
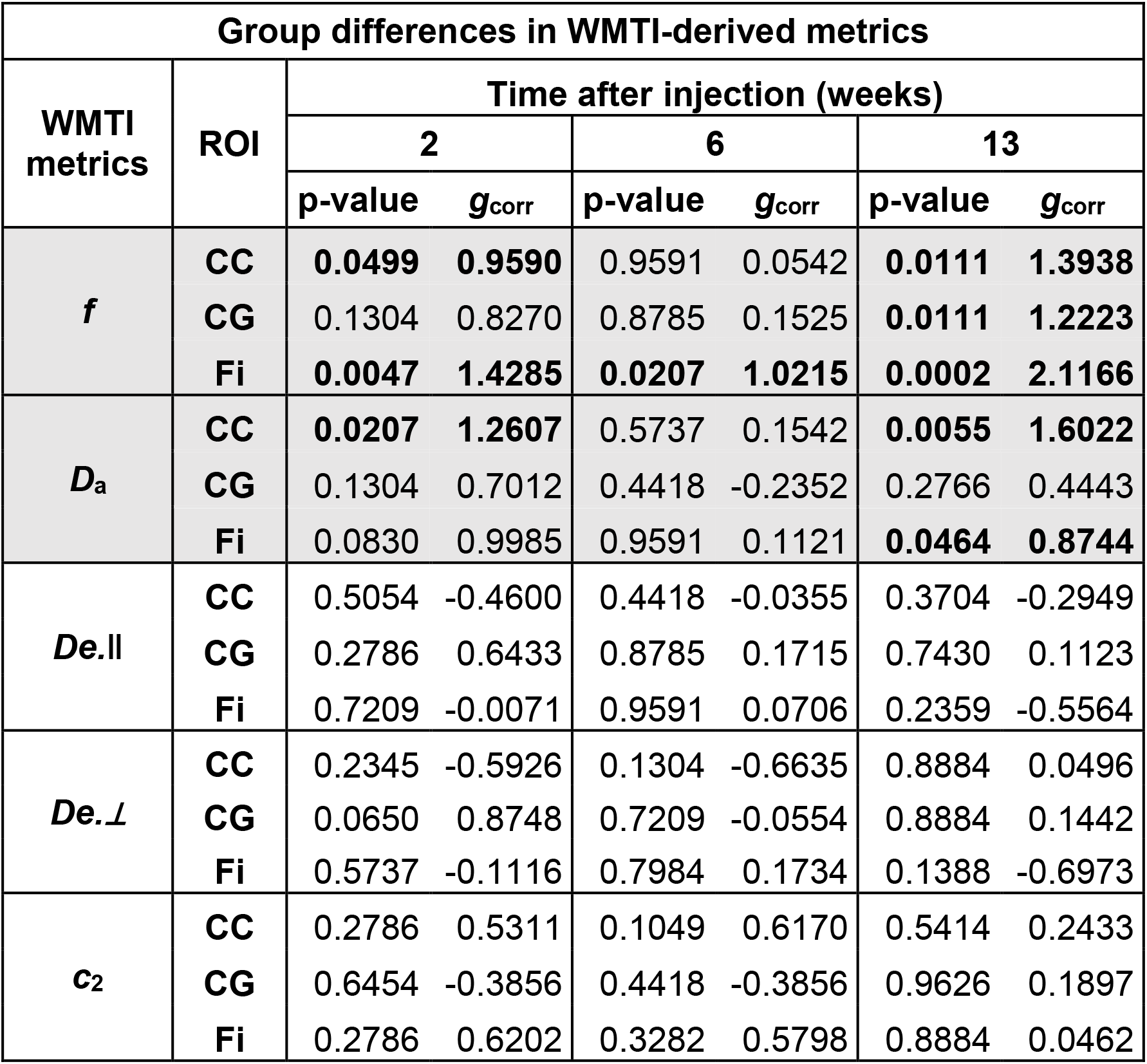
List of p-values obtained from the two-tailed Mann-Whitney *U*-test (α=0.05) and corrected Hedge’s g (g_corr_) used to evaluate differences in WMTI model metrics between STZ and CTL groups at each timepoint in each ROI. ROI: Region of interest; CC: corpus callosum; CG: cingulum; Fi: fimbria; f: volume fraction of water molecules; D_a_: intra-axonal diffusivity; D_e,║_: parallel extra-axonal diffusivity; D_e,⊥_: perpendicular extra-axonal diffusivity; *C*_2_: orientation dispersion. *Bold font: p < 0.05. Gray cells: ROIs shown in Figure 7.*

## Footnotes

1 https://github.com/NYU-DiffusionMRI/mppca_denoise

2 https://bitbucket.org/reisert/unring/

3 https://github.com/NYU-DiffusionMRI/DESIGNER

## Notes

### Competing Interest Statement

The authors have declared no competing interest.

